# Distinct effects of host and neighbour tree identity on arbuscular and ectomycorrhizal fungi along a tree diversity gradient

**DOI:** 10.1101/2021.05.18.444754

**Authors:** Olga Ferlian, Kezia Goldmann, Nico Eisenhauer, Mika T. Tarkka, François Buscot, Anna Heintz-Buschart

## Abstract

Plant diversity and plant-related ecosystem functions have been in focus in biodiversity-ecosystem functioning studies. However, in this context, biotic interactions with mycorrhizal fungi have been understudied although they are crucial for plant-resource acquisition. We investigated the effects of tree species richness, tree mycorrhizal type on arbuscular (AMF) and ectomycorrhizal fungal (EMF) communities. We aimed to understand how dissimilarities in taxa composition and beta-diversity are related to target trees and neighbours of the same/different mycorrhizal type. We sampled a tree experiment with saplings (∼7 years old), where tree species richness (monocultures, 2-species, and 4-species mixtures) and mycorrhizal type were manipulated. AMF and EMF richness significantly increased with increasing tree species richness. AMF richness of mixture plots resembled that of the sum of the respective monocultures, whereas EMF richness of mixture plots was lower compared to the sum of the respective monocultures. Specialisation scores revealed significantly more specialised AMF than EMF suggesting that, in contrast to previous studies, AMF were more specialised, whereas EMF were not. We further found that AMF communities were little driven by the surrounding trees, whereas EMF communities were. Our study revealed the drivers of mycorrhizal fungal communities and further highlights the distinct strategies of AMF and EMF.

## Introduction

Ecological research in the last decades has provided compelling evidence that biodiversity change alters ecosystem functioning [1, 2]. This relationship has been studied extensively in experiments and in natural ecosystems [2, 3]. Typically, plant diversity as well as plant-related ecosystem functions have predominantly been in the focus [4, 5], while cascading effects of biodiversity change at one trophic level to other trophic levels have attracted less attention [6, 7]. However, in this context, biotic interactions with plant symbionts, such as mycorrhizal fungi, remain unstudied, although they may be directly linked to plant-resource acquisition and, consequently, to plant competition and coexistence in plant communities [8–10]. Moreover, there is still a lack of knowledge of the factors that, besides plant diversity itself, influence plant-symbiont interactions and the impact of different forms of interactions on each other. One reason could be the still difficult assessment of plant symbionts that are often of microscopic size [11]. But due to the development of molecular tools [12], the access to soil-borne organisms has become easier.

The majority of plants are associated with a form of mycorrhiza, a close mutualistic interaction between roots and fungal partners [13]. The two partners are in cellular contact for nutrient exchange: the fungal partner receives photosynthetically fixed carbon, whereas the plant partner is supplied with nutrients, such as phosphorus and nitrogen [14]. Furthermore, the ability to tolerate abiotic and biotic environmental stresses increases in mycorrhizal plants [15, 16]. Mycorrhizal fungi are able to form large hyphal networks belowground that interconnect multiple host plants [17]. Such common mycorrhizal networks further contribute to the enhanced nutrient supply by gathering and sharing distant resources that are otherwise inaccessible to plant roots [13].

The two main groups of mycorrhizal symbioses on trees of temperate zones are arbuscular mycorrhiza (AM) and ectomycorrhiza (EM), which have distinct life strategies with respect to resource acquisition and allocation as well as interaction strength [13, 18, 19]. AMF can primarily access mineral nutrients, and only a few taxa are able to acquire P and N from organic sources [20, 21]. EMF have the ability to decompose dead organic matter [14, 22]. The general assumption that plants associate exclusively with one mycorrhizal type has been repealed by repeatedly detecting dual mycorrhization with AMF and EMF in plant roots [23, 24]. The extent of this dualism is often context-related and depends on environmental factors, such as nutrient availability, climatic conditions, or plant age [23]. A general prerequisite for the colonisation by specific fungal species as well as their diversity is the propagule reservoir in soil. Nevertheless, one of the two mycorrhizal types dominates the association with a plant host [25] with differential benefits from AMF and EMF colonisation, but there is limited information on the local drivers of dual mycorrhization and mycorrhizal diversity.

Higher plant diversity and, thus, more variance in root traits and microenvironments leads to a higher diversity of mycorrhizal fungi in experimental set-ups, where the diversity of plant communities was manipulated [8, 26, 27]. However, we lack knowledge on species composition of mycorrhizal fungi in diverse plant communities compared to those in their respective plant monocultures, i.e., whether compositions are additive or potentially follow other patterns, and on the respective underlying mechanisms. Furthermore, the characteristics of the associating fungal species, such as their specificity, may be of importance. Some mycorrhizal fungal species are shared by different plant species, whereas others are specialised on particular plants. Typically, AMF include more generalists that colonise several plant species than EMF [28, 29]. Also, AM and EM plants can have certain levels of specificity for particular mycorrhizal fungi and their specific characteristics [30, 31]. But to date, only few studies have been carried out in forest systems to explore tree neighbourhood effects on mycorrhizal diversity and community composition [32].

In our study, we took advantage of an existing diversity experiment with tree saplings (6.5 to 7.5 years old), where tree species richness (monocultures, 2-species, and 4-species mixtures) and root mycorrhization were manipulated via suitable tree species selection [33]. Tree communities of only AM, only EM, and of trees with both mycorrhizal types were set up. Our study represents a follow-up study to Heklau et al. [24] who analysed the effects of differently mycorrhized tree neighbours on target trees’ fungal community compositions at an earlier time point. We further studied the effects of tree species richness, tree mycorrhizal type, neighbourhood, and their interactions on AMF and EMF specialisation, community richness, phylogenetic diversity, and beta-diversity. We hypothesized that (1) AMF richness of all tree species in tree species mixtures is lower than the sum of the respective number of AMF species of the tree species in monocultures and that EMF display the opposite pattern. Hypothesised mechanisms are the comparably specialist strategy of EMF, the generalistic strategy of AMF, as well as the selection for generalistic AMF being increasingly shared by tree species in mixtures. We further hypothesised that (2) AM trees have a higher AMF richness and EM trees have a higher EMF richness, and, thus, the former comprises a higher proportion of specialised AMF and the latter a higher proportion of specialised EMF. (3) Due to the dominance of generalistic species in AMF, mycorrhizal associations are expected to be driven by tree species identity and/or mycorrhizal type of the tree neighbour in the case of AMF but not of EMF. Thus, AMF composition is hypothesised to be more similar among neighbouring tree species than EMF composition.

## Results

### Sequencing success and mycorrhizal fungal richness

All roots were checked for mycorrhization, and the colonisation rates were microscopically determined for AMF and EMF (Supplementary Table S1). AM colonisation rates were higher in AM trees (1.58 - 25.81%) than in EM trees (0.85 - 2.06%). EM colonisation rates were higher in EM trees (59.49 - 88.33%) than in AM trees (0.00 - 1.07%), where they were equal to zero in four of the five tree species.

The sequencing and the subsequent bioinformatic analyses led to 62 AMF VT in 179 root samples, and 174 EMF ASVs in 152 root samples.

Overall, the mean AMF richness per plot was slightly higher than the one for EMF (Supplementary Table S2). However, the fungal richness varied according to the tree mycorrhizal type of the trees, i.e. there were more AMF VT associated with roots of AM trees and more EMF ASVs in EM roots. Furthermore, plots with only AM trees had a higher AMF richness compared to plots with only EM trees. Plots with both mycorrhizal types had an intermediate AMF richness. For both AMF and EMF, fungal richness increased significantly from tree monocultures to mixtures (ANOVA, P < 0.05).

### Expected vs. observed fungal richness

The correlation between observed (number of unique fungal species detected in all trees of a mixture plot) and expected (number of unique fungal species in the respective monocultures of the tree species in mixture plots) AMF richness was positive in all treatments, except for EM plots with four species (Fig. 1a). However, only the correlations of AMF richness in AM plots with two tree species and those in plots with both mycorrhizal types with two tree species were significant (Table 1). Regression models of four-species mixtures had generally lower slopes and deviated more from the 1:1 line than models of two-species mixtures, indicating that the observed fungal richness decreased relative to the expected richness with higher tree species richness.

**Fig. 1.**
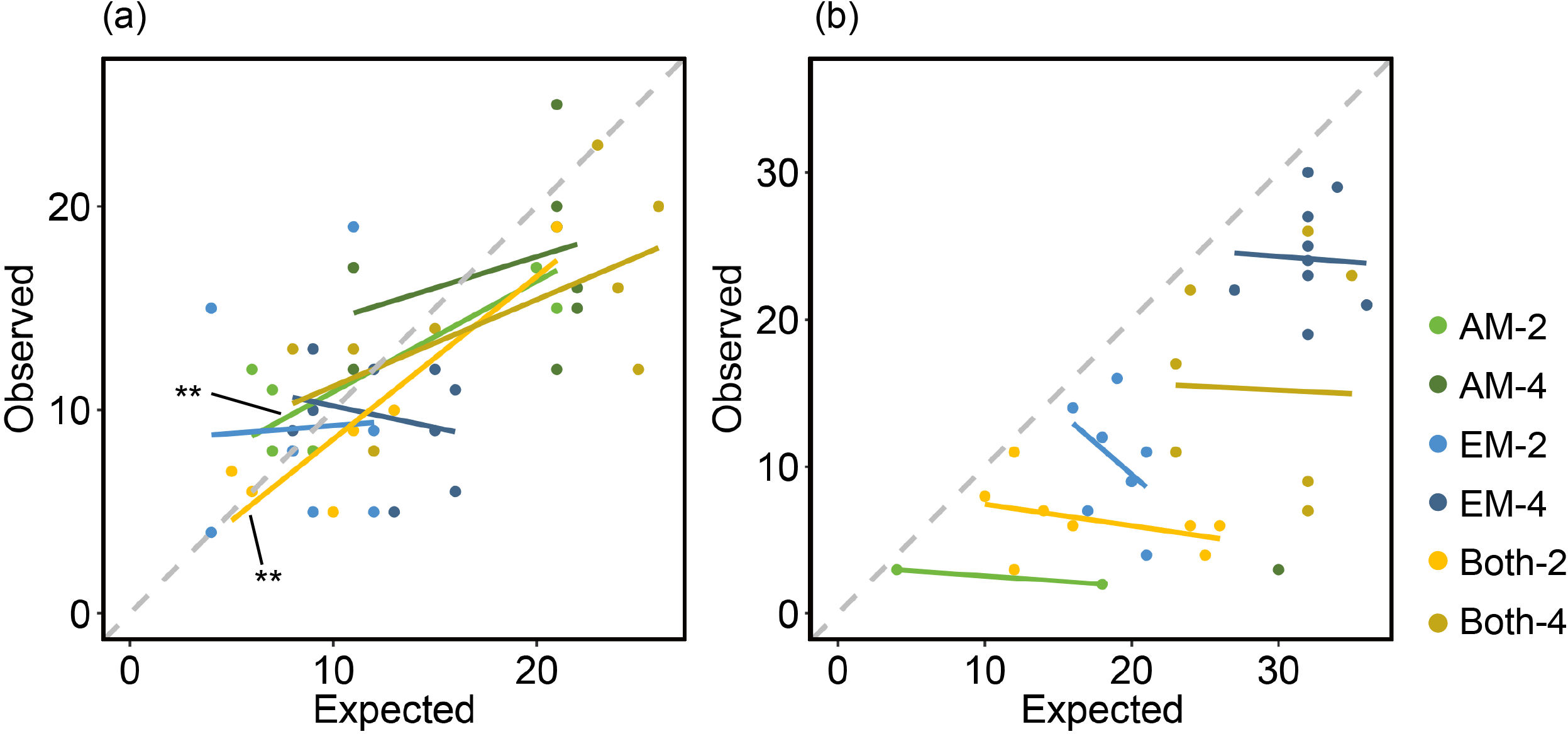
Correlations between expected and observed unique (a) arbuscular mycorrhizal fungal and (b) ectomycorrhizal fungal richness in plots with AM, EM, and both mycorrhizal type trees (AM, EM, and Both), in 2- and 4-species mixtures (−2 and -4). The grey dashed lines represent the 1:1 line, where expected equals observed fungal richness. Treatments whose data did not allow for statistical analyses (low detection rates) do not have a regression line. Asterisks indicate significant correlations (**P < 0.01) (N = 188).

**Table 1.** Summary of correlation analyses (Pearson’s correlation coefficient) between expected and observed richness of arbuscular mycorrhizal fungi (AMF) and ectomycorrhizal fungi (EMF) in mixture plots, respectively. Analyses were conducted separately for the six experimental treatments (mycorrhizal type of tree: AM, EM, and Both; tree species richness levels: 2 and 4). Due to low detection rates of EMF-ASVs, two of the treatments could not be statistically tested. Significant relationships are highlighted in bold (P < 0.05) (N = 188).

|  | AMF richness |  |  | EMF richness |  |  |
| --- | --- | --- | --- | --- | --- | --- |
|  | df | r | P | df | r | p |
| AM-2 | **6** | **0.86** | **0.01** | - | - | - |
| AM-4 | 7 | 0.34 | 0.37 | - | - | - |
| EM-2 | 6 | 0.05 | 0.91 | 7 | -0.42 | 0.26 |
| EM-4 | 7 | -0.25 | 0.52 | 8 | -0.05 | 0.88 |
| Both-2 | **5** | **0.91** | **0.00** | 6 | -0.39 | 0.34 |
| Both-4 | 7 | 0.63 | 0.07 | 6 | -0.03 | 0.94 |

In contrast to the patterns in AMF, all correlations between expected and observed EMF richness had negative trends (Fig. 1b). However, no single correlation was significant (Table 1). Correlations could not be analysed in plots with only AM trees due to the low number of samples where EMF-ASVs were detected. Moreover, all data points were distributed below the 1:1 line, indicating lower than expected EMF richness in the mixture plots.

Overall, the AMF data points were more evenly distributed around the 1:1 line, and regression lines were relatively similar to the 1:1 line in terms of slope and position than the EMF data points. Consequently, divergence between AMF expected and observed fungal richness was overall lower than that for EMF with lower EMF fungal richness in the mixtures than expected.

### Taxonomic overview

The taxonomic assignments gained from the fungal amplicon sequencing were supported by results retrieved from Sanger sequencing of mycorrhizal structures after the morphotyping (Supplementary Results S1, Table S3). Although not all fungal ASVs were found with this traditional sequencing method, we showed that the high-throughput approach covered the living fungi.

The AMF belonged to eight different genera, with a majority belonging to *Glomus* (35 AMF). Only three AMF appeared in all studied treatments (Fig. 2a). The overall AMF richness increased with increasing AM tree species richness, but decreased again if there was an EM tree present (ANOVA, P < 0.05; Fig. 2a). EM trees also had AMF. In fact, even there, the AMF richness increased from EM tree monocultures to mixtures (ANOVA, P < 0.05; Fig. 2a). However, the highest richness coupled with the biggest variety of AMF genera in EM tree treatments was found in four species mixtures, where EM and AM trees occured together (Fig. 2a). Only three AMF taxa were found in all treatments (Fig. 2a). In contrast, the EMF belonged to 16 different genera (seven Ascomycota and nine Basidiomycota genera) and no EMF appeared in all treatments (Fig. 2b). The overall richness of Ascomycota was higher in AM tree treatments and Basidiomycota predominated when an EM tree species was also present (ANOVA, P < 0.05; Fig. 2b). The overall EMF richness was considerably lower in AM trees, but a general increase of EMF richness with increasing AM tree diversity level was detectable (ANOVA, P < 0.05; Fig. 2b). Likewise, the overall EMF richness linearly increased with increasing diversity of EM trees and, again, decreased in the treatments where AM and EM trees were mixed (Fig. 2b). Generally, we found a higher proportion of AMF taxa shared by multiple experimental treatments compared to EMF taxa, and the shared AMF taxa were also present in larger sets of treatments in AMF compared to EMF.

**Fig. 2.**
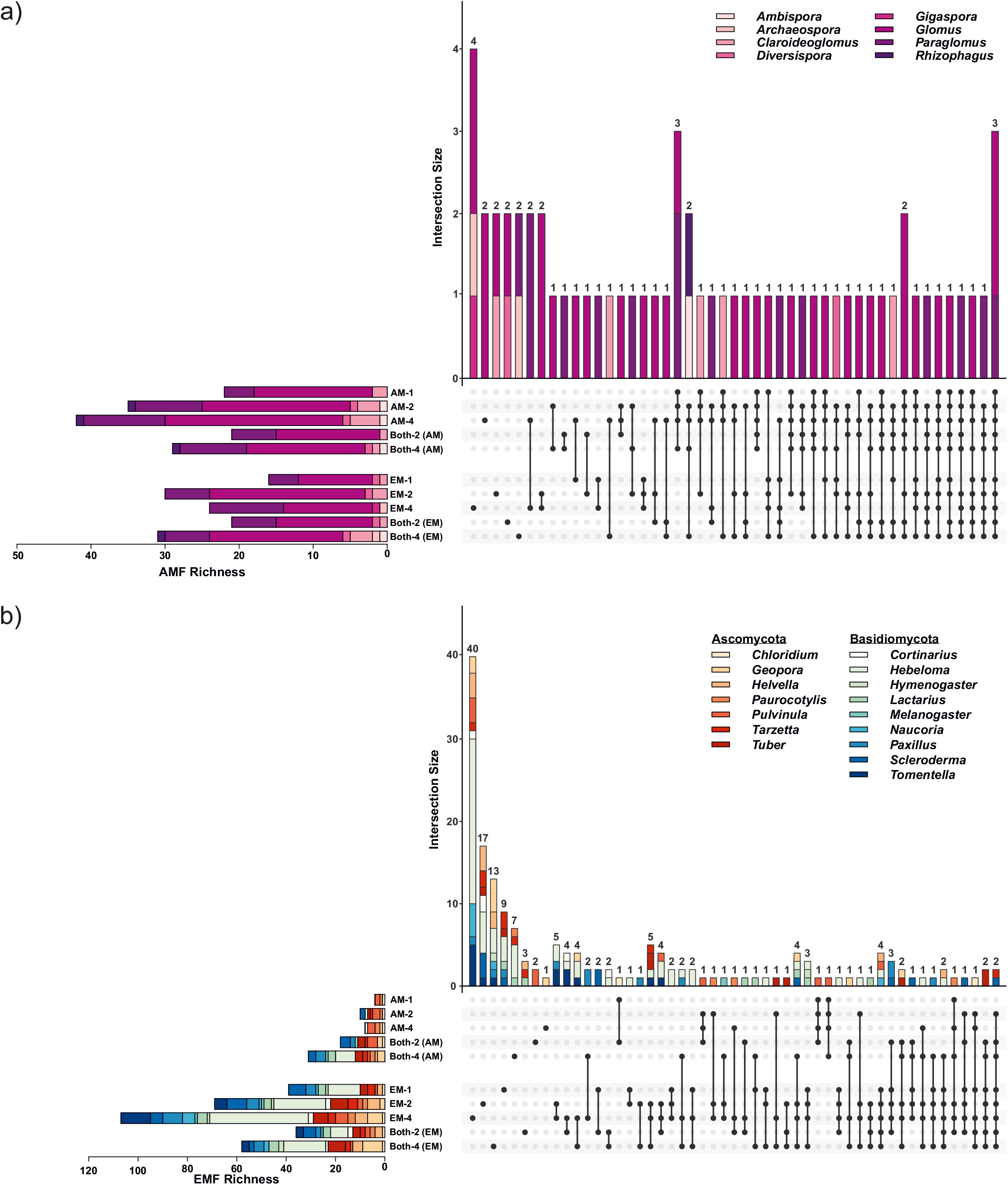
UpSet plots displaying for a) AMF and b) EMF, the richness and composition of the treatments overall (horizontal bars) and of specific and shared subsets of fungal taxa (vertical bars) according to the intersection matrix. Connected dots represent intersections of shared fungal taxa. Horizontal bars of each panel represent the mycorrhizal fungal richness as absolute count values, colour-coded according to the assigned fungal genera. Vertical bars show the intersection size between fungal communities in the different treatments. Numbers above vertical bars represent the number of fungi taxa found in the treatment(s) marked by the black dot(s) (N = 188).

### Specialisation of fungi

We calculated a score to evaluate the specialisation of mycorrhizal fungi along the gradient of tree species richness (Fig. 3). This revealed a constant, high specialisation of AMF to AM and EM trees in tree monocultures and mixtures, with the exception of the treatments with only EM trees (ANOVA, P < 0.05). Likewise, the EMF specialisation was high in all treatments except the ones containing only AM trees (ANOVA, P < 0.05).

**Fig. 3.**
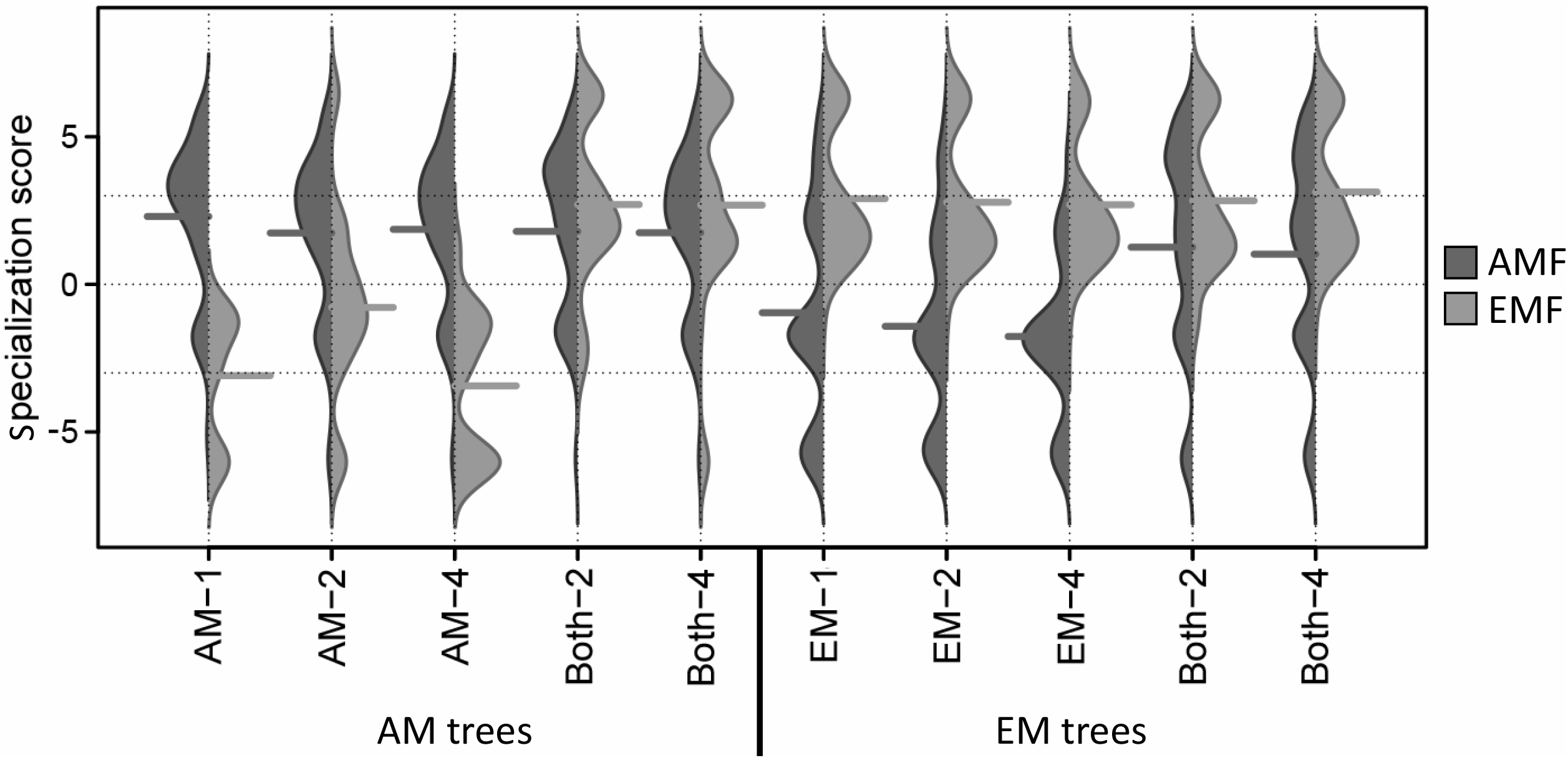
Specialisation scores of arbuscular mycorrhizal fungi (AMF) virtual taxa and ectomycorrhizal fungi (EMF) amplicon sequencing variants in trees with predominantly arbuscular mycorrhiza (AM trees) and trees with predominantly ectomycorrhiza (EM trees) across all treatments, respectively (mycorrhizal type of tree: AM, EM, and Both; tree species richness levels: 1, 2 and 4). The score represents the difference between the specialisation phi of a fungal taxon to its respective tree type and the mean of the specialisation of 100 null models for this taxon, divided by the standard deviation of the 100 null models for this ASV. The dashed lines indicate 0 and 3 standard deviations (N = 188).

The degree of average specialisation was higher in AMF than EMF (Wilcoxon rank sum test with continuity correction, P < 0.001). Thereby, the AMF and EMF found in tree species mixtures appeared to be rather associated to AM and EM trees, respectively. This means that trees on plots with both mycorrhizal types have EM tree-specific EMF and only very limited AM tree-specific EMF and *vice versa*.

This analysis also allowed the identification of specific AMF or EMF taxa that were highly specialized to trees of either mycorrhizal type (Supplementary Table S4). The findings revealed that genera like *Glomus* and *Paraglomus*, which include the majority of AMF, showed high specialisation. Among those, the number of specialised AMF on AM trees was higher than on EM trees. Similarly, more EMF were specialised on EM trees than on AM trees.

### Tree host species and neighbour effects on phylogenetic diversity

AMF phylogenetic diversity in AM trees was significantly affected by tree species identity of the target tree and, in EM trees, by mycorrhizal type of the tree neighbours (Table 2). AMF phylogenetic diversity in EM trees was significantly higher when the tree neighbour was of the other mycorrhizal type compared to a neighbour of the same type (Fig. 4a). EMF phylogenetic diversity in AM and EM trees was significantly affected by mycorrhizal type of the tree neighbours and, in EM trees, further by tree species identity of the target tree (Table 2). EMF phylogenetic diversity in AM trees was significantly higher when the tree neighbour was of the other mycorrhizal type compared to a neighbour of the same type (Fig. 4b). In contrast, in EM trees, EMF phylogenetic diversity was significantly lower when the tree neighbour was of the other mycorrhizal type compared to a neighbour of the same type. AMF and EMF phylogenetic diversities did not differ between the other treatments.

**Table 2.** Summary of linear effects analyses on phylogenetic diversity of arbuscular mycorrhizal fungi (AMF PD) and ectomycorrhizal fungi (EMF PD) as affected by tree species identity of the target tree, tree species of the neighbour tree (same vs. different), mycorrhizal type of the neighbour tree (same vs. different), and tree species richness of the plot. Analyses were conducted separately for trees predominantly associated with arbuscular mycorrhizal fungi (AM trees) and trees predominantly associated with ectomycorrhizal fungi (EM trees). Significant effects are highlighted in bold (P < 0.05) (N = 188).

|  |  | AMF PD |  | EMF PD |  |
| --- | --- | --- | --- | --- | --- |
|  | df | X<sup>2</sup> | P | X<sup>2</sup> | P |
| **AM trees** |  |  |  |  |  |
| Tree species target | 4 | **69.70** | **< 0.001** | 2.55 | 0.64 |
| Tree species neighbour | 1 | 2.63 | 0.10 | 0.29 | 0.59 |
| Mycorrhizal type neighbour | 1 | 0.08 | 0.78 | **36.94** | **< 0.001** |
| Tree species richness | 1 | 0.01 | 0.94 | 0.05 | 0.83 |
| **EM trees** |  |  |  |  |  |
| Tree species target | 4 | 5.22 | 0.27 | **10.84** | **0.03** |
| Tree species neighbour | 1 | 0.15 | 0.70 | 2.01 | 0.16 |
| Mycorrhizal type neighbour | 1 | **7.49** | **0.01** | **17.48** | **< 0.001** |
| Tree species richness | 1 | 0.10 | 0.76 | 0.13 | 0.72 |

**Fig. 4.**
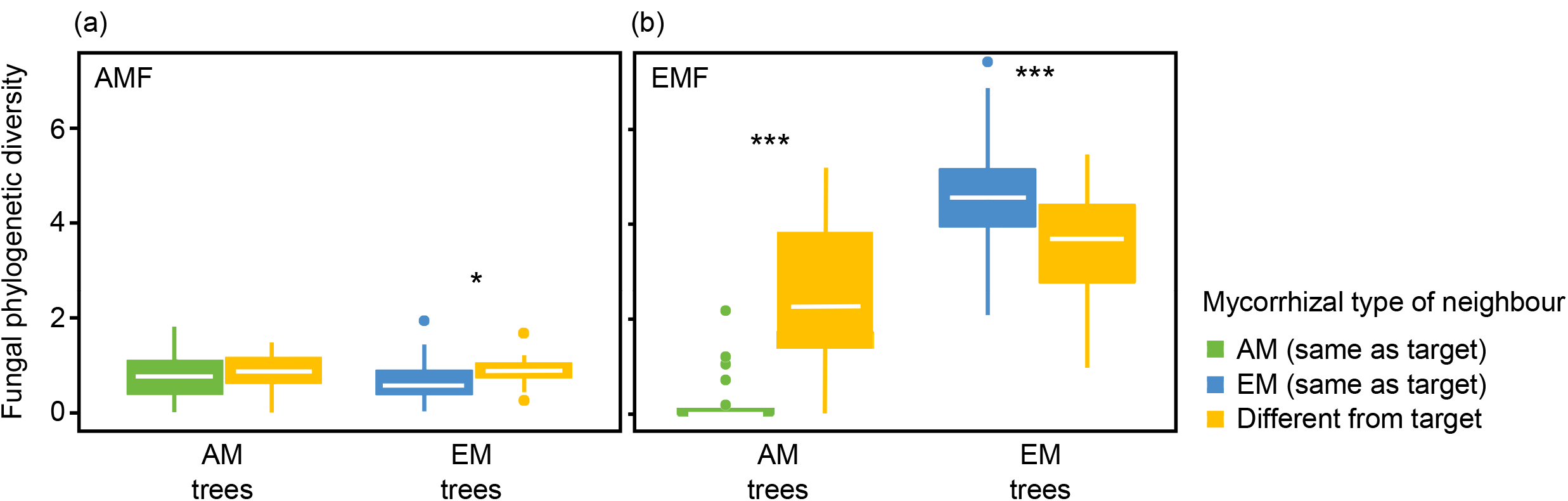
Fungal phylogenetic diversity of (a) arbuscular mycorrhizal (AMF) and (b) ectomycorrhizal fungi (EMF) in roots of target trees predominantly forming arbuscular mycorrhiza (AM trees) and that forming ectomycorrhiza (EM tress) as affected by the mycorrhizal type of the tree neighbour. Significant differences are indicated by asterisks. *P < 0.5, ***P < 0.001 (N = 188).

### Similarities of mycorrhizal fungal communities among tree species and communities

To understand if the fungal communities associated with trees in mixtures are more similar to the respective monoculture communities or to the communities of the tree neighbours, we analysed the pairwise Soerensen similarities of the mycorrhizal fungal communities (Fig. 5). AMF communities in mixtures were more similar to monocultures of the same tree species than to their neighbours within mixtures, indicating tree species-specific communities (Fig. 5a, b). In comparison, EMF communities of target trees were more similar to those of tree neighbours than between the target tree in monocultures and mixtures indicating mixture-adapted communities (Fig. 5c, d). This pattern was stable when comparing monocultures to two or monocultures to four tree-species mixtures.

**Fig. 5.**
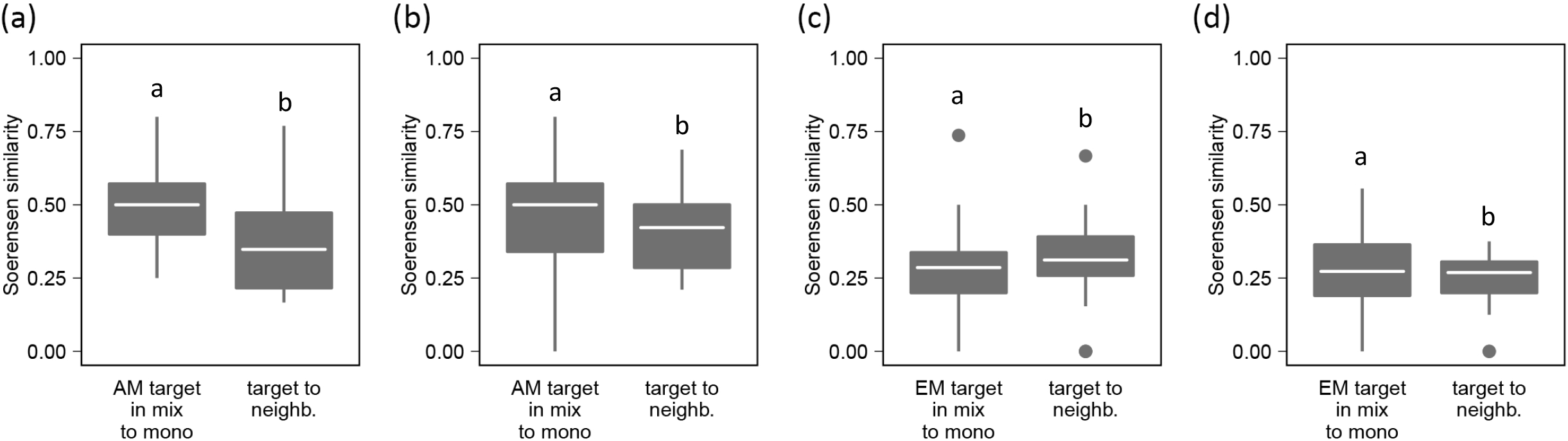
Comparisons of pairwise beta-diversities. The left box of each panel represents the Soerensen similarity between mycorrhizal fungal communities of target tree in mixture and that of monoculture; the right box represents the Soerensen similarity between mycorrhizal fungal communities of the target tree and that of their the neighbouring tree; a) AMF communities of AM trees in two species mixtures (AM-2); b) AMF communities of AM trees in four species mixtures (AM-4); c) EMF communities of EM trees in two species mixtures (EM-2); d) EMF communities of EM trees in four species mixtures (EM-4). Letters above boxplots indicate significant differences according to Wilcoxon rank sum tests, P < 0.05 (N = 188).

## Discussion

Overall, our study showed that AM colonisation rates were relatively low in AM tress and even lower in EM trees; EM colonisation rates were higher in EM trees than in AM trees. With increasing tree species richness, AMF as well as EMF richness increased. Surprisingly, AMF richness of mixture plots resembled that of the sum of the respective monocultures, whereas EMF richness of mixture plots was lower compared to the sum of the respective monocultures. Zooming into this pattern, we found that tree species in mixtures more commonly shared EMF than AMF, suggesting that EMF tended to be more generalistic than AMF in our study. This was supported by the finding that EMF diversity and composition of target trees were more strongly influenced by the mycorrhizal identity of the tree neighbour in comparison to AMF.

### Fungal richness in AM vs. EM trees

We found expected differences in the mycorrhizal community compositions between AM and EM host trees with more AMF than EMF on AM trees and more EMF than AMF on EM trees. Morphologically assessed mycorrhizal colonisation rates of AM and EM supported these findings. This is in line with findings in Heklau et al. [24], where root mycorrhizal communities were characterised using morphological and next-generation sequencing techniques in the same experiment two years prior to our assessments. Interestingly, both studies showed that all tree species had a dual mycorrhization with AM and EM trees being more equally colonised by AMF than by EMF. This suggests that the two tree species groups (AM and EM trees) rather form two distant tree species pools along a continuum of AMF-to-EMF colonisation instead of two distinct characteristics.

Interestingly, typical forest EMF taxa, such as *Russula* or *Inocybe* [34], were not found on our sampled trees which, in contrast, were rich in members of Pezizales that have been associated with young seedlings during early phases of forestation [35, 36]. Possible reasons could be that the site used to be an agricultural field without any forest stands in the surroundings before setting up the plots [37]. Accordingly, the reservoir for EMF propagules can be estimated as poor, most likely with only a small potential for pioneer EMF at the beginning of the experiment [38]. Moreover, even after the trees were planted, the coverage with the weed tarp to minimise weed interference could have led to a decreased amount of propagules entering the soil and tree roots via wind or animal dispersal. In addition, the form and extent of mycorrhizal associations are typically context-dependent, e.g., dependent on abiotic factors, such as nutrient availability [23, 39]. The specific nutrient- and humus-rich Chernozem soil of the MyDiv experimental site could have restricted common EMF infections that typically supply the plant host with nutrients from the decomposition of dead organic matter.

Overall, among the AMF, we predominantly found members of the Glomerales, such as *Glomus*, that are characterised as r-strategists, i.e. with a short generation time, fast growing hyphae, and low resource use efficiency [40, 41]. Their ability to adapt and reproduce quickly may have led to this *Glomus* dominance that in turn may have facilitated the dominance of rather generalistic AMF, and they have been reported to occur in high abundances in forests before by Öpik et al. [42]. The dominance by *Glomus* could lead to an outcompeting of other AMF and, accordingly, cause a decrease of AMF richness [43]. However, we did not observe a decrease in AMF richness, but rather a high species diversity within the genus *Glomus*. Besides, the *Glomus* dominance may partially be due to an over-representation of sequences affiliated to this genus by the amplification of the SSU marker region [41].

Our results further showed that both AMF and EMF richness of the target tree were positively related to tree species richness of the plot, which is in line with previous studies [44, 45]. The observations support the idea of a sampling effect, meaning that in more diverse plant communities, the probability of having species with distinct abilities to associate with particular fungal species increases [46]. Higher plant diversity also facilitates diverse microenvironments and divergent niches for soil microorganisms in general [8, 26, 27]. In addition, higher plant diversity is well-known to increase plant biomass and, consequently, the extent and diversity of carbon inputs into the rhizosphere [47] facilitating the coexistence of a multitude of fungal species [48]. However, the majority of previous studies exploring the plant diversity–fungal diversity relationship assessed fungal communities in soil (plot level) rather than in host tree individuals, although the latter is more indicative of the actual interactions between fungi and plants [26, 46] (but see [8, 44]). Saks et al. [49] found that AMF community composition in soil was rather random and influenced by environmental factors compared to AMF in roots, where also comparably more species were found. Given the fact that these relationships hold true for communities in both soil and tree roots, we can presume that one important factor limiting root colonisation of particular fungal species is its availability in soil.

### Specialisation in mycorrhizal fungi

Overall, AMF richness of all tree species (in all treatments) in mixtures resembled that of the sum of their respective monocultures. In contrast, EMF richness of all tree species in mixtures tended to be generally lower than the sum of their respective monocultures. This suggests that the AMF assemblages in mixtures were composed of distinct unique fungal species that specifically associate with particular tree species, irrespective of the surrounding plant community composition. In contrast, the sum of unique EMF increased comparably little from monocultures (expected) to mixtures (observed), indicating that tree species in mixtures share part of the EMF species. These findings are exactly opposite to what we hypothesised. Interestingly, in four-species mixtures, we found a slight but consistent weaker relationship between expected and observed richness compared to two-species mixtures, both for AMF and EMF. This shows that with increasing tree diversity (from two to four), mycorrhizal fungal species associated with respective tree species increasingly overlap among the tree species. Tree species mixtures, in contrast to monocultures, contain more potential host plants and create a relatively heterogeneous soil environment in terms of resources, which may consequently, favour generalistic over specialised fungal species [8], explaining the increasing proportion of generalistic fungi with increasing plant diversity.

Calculations of specialisation coefficients revealed that AM and EM trees associated preferably with specialised AMF and EMF, respectively, which confirmed our second hypothesis. The results suggest that the mycorrhizal fungi belonging to the opposite mycorrhizal type than the host tree are comparably rare and rather generalistic taxa. This was further supported by the finding that, in AM trees, the species identity of the target tree only drove the phylogenetic diversity of the fungal community belonging to the same mycorrhizal type. This may partly be due to the marginal EM colonisation of AM trees. Molina and Horton [50] reviewed plant host preferences of AMF and EMF and argued that hosts may preferably allocate resources to specialised rather than generalistic fungi, as they benefit more from interactions with specialist fungi. Within the two major EMF phyla, Ascomycota were more dominant than Basidiomycota on AM trees in AM treatments, whereas Basidiomycota predominated on AM trees when EM trees were present, which suggests different specificities of the EMF of the two phyla. In AM trees, for example, there were far more AMF than EMF which points to a particular selectivity that may lead to a high proportion of specialised AMF in AM trees compared to EM trees.

In general, we found more specialised AMF than EMF confirming the idea of a more specialised strategy in AMF and a more generalistic strategy in EMF (EMF sharing [39]), which may also mechanistically explain the observed patterns of the relationships between expected and observed fungal species richness. van der Linde [51] also reported that only approximately 10% of EMF are host-specific. However, this is in contrast to our main hypothesis that is based on the theory of the fungi’s evolutionary history and previous findings, where AMF are commonly assumed to be generalists, whereas EMF are host-specific [49, 52]. Generalistic AMF were also confirmed by Weißbecker et al. [8], who used the same measure of species specialisation, but analysed soil from the root zone. Specialisation estimates mostly originate from lab studies or specific contexts. Most cultivable AMF species used in lab experiments are generalists [50]. Contexts, such as ecosystem (e.g., grasslands/agricultural sites vs. forests), biome (e.g., (sub)tropical vs. temperate), and host type (e.g., gymno-vs. angiosperms) may drive the distribution of AM and EM [53], and it, thus, seems likely that they drive their specificity. In any case, it has to be noted that in both, AMF and EMF, specialised and generalistic species exist [50].

Specialisation scores further revealed that adding tree species of a different mycorrhizal type to a plant community resulted in an increase of specialised mycorrhizal fungi in the target tree when the added tree was of the same mycorrhizal type as the fungal type in question. Some mycorrhizal fungi seemed to depend on tree communities, where AM and EM trees grew together, which may point to a facilitation by neighbouring trees or the existence of mycorrhizal fungal species that form common mycorrhizal networks among trees of different mycorrhizal types [50, 54]. As most mycorrhizal trees were found to have dual mycorrhization, it is not surprising that AM and EM trees may also interconnect. In our study, this effect could be detected for AMF as well for EMF. Such potential transfer of resources may considerably complement other acquisition strategies of trees and contribute to an increase in ecosystem functioning in communities with trees of different dominant mycorrhizal types [33].

### Effects of tree neighbour identity

Our results indicated that fungal diversity was driven by tree species identity of the target tree and the mycorrhizal type of the tree neighbour. Surprisingly, tree species identity of the neighbour or tree species richness of the community did not significantly affect fungal diversity. These results applied to both AMF and EMF but were found to be more pronounced in EMF, which refutes our third hypothesis assuming that exclusively AMF communities are affected by the identity of the tree neighbour. This suggests that beyond the local availability of infective propagules, such as spores and hyphae, in soil [49], two important factors determining mycorrhizal associations (host plant and biotic environment [55]) are underpinned by distinct mechanisms. This is valid for different mycorrhizal types. Associations with trees depended on host species identity; however, the effect of the neighbouring tree community was rather driven by their mycorrhizal identity. This is in line with several previous studies (e.g., Dickie et al. [56]), but contrasts more recent findings that fungal specificity on host plants, especially in AMF, is targeted at a broader unit, such as the ecological-group level of the plant species [42, 57].

We found that fungal AMF diversity on EM trees was increased by the presence of AM tree neighbours, and EMF diversity on AM trees was increased by the presence of EM tree neighbours but not that of the other fungal group, respectively. Although all tree species have a dual mycorrhization with both mycorrhizal types [23], the mycorrhizal fungi species within each fungal group seem to be distinct between AM and EM trees. This may be an effect of the opposing dominance of the mycorrhizal types in AM and EM trees. Such specific tree neighbours may facilitate mycorrhizal associations of a target tree. A neighbour effect was suggested, where the litter produced by tree neighbours may trigger specific fungal communities that colonise the target tree [50]. This hypothesis may have limited relevance in our study, as all of our plots were covered with tarp that prevented leaf litter material, the dominant form of litter in our system, from entering the soil. Moreover, the mycorrhizal identity of the tree neighbour may (more than tree species) influence the nutrient availability for the target tree directly or indirectly via altering soil microbial communities and, thus, its associations with fungal partners it exchanges resources with (Singavarapu et al. under review). Another potential interaction between target and neighbour trees are common mycorrhizal networks that need a specific tree partner of AMF and EMF [23].

For EMF diversity, we further found that EM trees were not only unaffected by the presence of AM tree neighbours on the plots, but EMF diversity even decreased, which may point to a dilution effect, as stated in Heklau et al. [24]. It describes that the presence of an unfavourable tree in the surrounding (in our case a tree of the other mycorrhizal type) may dilute potential interaction partners and, thus, limit the mycorrhizal association of the target tree.

Using the Soerensen similarity index, we also found that AMF communities of a specific tree species were more similar between respective individuals in monocultures vs. mixtures than between individuals of different tree species within a plot indicating a species-specific community (in both 2-species and 4-species mixtures). EMF communities showed the opposite pattern indicating a mixture-adapted community. This further supports our findings stated above that AMF communities were poorly predicted by the neighbouring plant community and, thus, comparably specialised, whereas EMF communities were comparably generalistic.

## Conclusions

Our study identified factors influencing the diversity of AMF and EMF communities associated with deciduous tree roots in a temperate forest plantation. Hereby, tree diversity, host species identity, and the mycorrhizal type of the surrounding plant community played significant roles. Building on the results found in Heklau et al. [24], we were able to further identify factors predicting mycorrhizal fungal communities and explaining the lack of additive effects of fungal species from tree monocultures to mixtures. We found substantial differences in the specificity of AMF and EMF. In contrast to our expectations and previous studies, AMF showed a more specialised and EMF a more generalistic strategy. The unexpected result highlights the fact that few studies have explored fungal communities of both AMF and EMF in parallel in temperate deciduous tree species. Furthermore, our study sheds light on the dual mycorrhization of trees which has been documented repeatedly but not explored in terms of the ecological specifics of the fungal partners that were considerably different between AM and EM trees. Finally, the type of the surrounding plant community played a more important role for EMF than for AMF, pointing to the importance of thorough plant species selection in forestry, e.g., for the set-up of productive tree plantations. Such insights help understanding the role of plant symbionts in plant interactions and competition, and, consequently, in the general mechanisms underlying biodiversity-ecosystem functioning relationships.

## Material and Methods

### Study site

The study was carried out as part of the MyDiv Experiment, a tree diversity experiment located in Saxony-Anhalt, Germany, at the Bad Lauchstädt Experimental Research Station of the Helmholtz Centre for Environmental Research – UFZ (51°23’ N, 11°53’ E [33]). The elevation is 115 m a.s.l.; the continental climate has an annual mean temperature of 8.8°C and a mean annual precipitation of 484 mm; the parent material is silt over calcareous silt; the soil type is Haplic Chernozem developed from Loess with a pH ranging between 6.6 and 7.4 [33, 58]. For more detailed site characteristics, see Ferlian et al. [33].

The MyDiv site encompasses 80 11×11 m plots that were set up in March 2015, with a core area in the centre of each plot (8×8 m). Each plot contains 140 trees in a 1 m-planting distance. All plots were covered by a water-permeable weed tarp to minimise weed interference. A total of ten tree species, five AM and five EM trees, were either planted in replicated monocultures, two-species or four-species mixtures [33]. Moreover, the design implemented a mycorrhizal type treatment represented by communities with only AM trees, only EM trees, or a combination of AM and EM trees in mixtures [33]. There was no direct control of mycorrhizal fungal association; and the treatment was established through assignment of tree species to dominant mycorrhizal types based on literature review (e.g., Wang and Qiu [59]) and respective planting. The following deciduous tree species were selected for the AM tree species pool: *Acer pseudoplatanus, Aesculus hippocastanum, Fraxinus excelsior, Prunus avium*, and *Sorbus aucuparia*; and the EM tree species pool: *Betula pendula, Carpinus betulus, Fagus sylvatica, Quercus petraea*, and *Tilia platyphyllos*. Per tree species, two monocultures were established. Furthermore, ten replicates per species richness level and mycorrhizal type were established, distributed over two blocks.

### Root sampling

Extending the sampling design of Heklau et al. [24], 200 root samples, one per plot and tree species, were taken in November 2019. In total, root samples from all 20 monocultures, 30 two-species mixtures, and 30 four-species mixtures were taken, ensuring that the correct individuals were sampled and avoiding contaminations (see Supplementary Methods S1).

### Quantification of mycorrhizal colonisation

AM colonisation of roots was quantified following Vierheilig et al. [60] by bleaching the roots in 10% KOH at 60°C overnight and staining the roots in a solution of 10% ink, 10% concentrated acetic acid, and 80% water. AM colonisation was quantified by assessing the abundances of arbuscules, vesicles, and hyphae with the gridline-intersect method [61]. The degree of EM colonisation was determined from fresh roots under a dissecting microscope accounting for differences in fine root morphology, colour, thickness, texture, and branching patterns of rootlets. From each sample, ten ∼5 cm root pieces of the first order were identified as colonised with EM when having a lighter colour and swollen tips, otherwise they were counted as inactive or not colonised. For analysis, frequency of EMF in percent was calculated as the proportion of root tips with active mycorrhiza in relation to all root tips examined.

### Identification of colonising mycorrhizal fungi via Sanger sequencing

For identification of root-inhabiting fungi, rootlets with ten EM root tips of EM trees or ten lateral roots of AM were harvested (from the samples for quantification of mycorrhizal colonisation) to polyethylene glycol 200 (Sigma-Aldrich, St. Louis, USA) adjusted to pH 13. The roots were extracted mechanically with glass beads by vortexing to release their DNA into the liquid. ITS regions were amplified according to White et al. [62] with the primers ITS1 (10 μM – 5′-TCCGTAGGTGAACCTGCGG) and ITS4 (10 μM – 5′-TCCTCCGCTTATTGATATGC), and Promega Green (Promega, Madison, USA) and sequenced with the ITS1 primer using Big Dye Termination Mix (GeneCust Europe, Dudelange, Luxembourg). Sequence quality was manually controlled using Sequencher 5.4.5. Sequences were compared to the UNITE database 8.0 using BLASTN 2.8.1. The raw Sanger sequences were deposited in the National Center for Biotechnology Information (NCBI) Genbank database under the accession numbers MW695221-MW695361.

### DNA extraction and Illumina sequencing

The root samples were first chopped and then manually ground using a porcelain mortar and liquid nitrogen. Approximately 0.2 g of the pulverised material was used for DNA-extraction using the Quick-DNA™ Fecal/Soil Microbe Miniprep Kit (Zymo Research Europe, Freiburg, Germany) following the manufacturer’s recommendations. Fungal ITS2 regions were amplified following the descriptions in Prada-Salcedo et al. [10] and AMF SSU regions were amplified following a nested PCR approach [63] (see Supplementary Methods S2). Libraries were prepared using Illumina Nextera XT and used for paired-end sequencing of 2×300 bp with a MiSeq Reagent kit v3 on an Illumina MiSeq platform. The raw Illumina sequences were deposited in the SRA of NCBI under the BioProject accession number PRJNA706719.

### Bioinformatics

Sequencing reads were processed to amplicon sequence variants (ASVs) using the DADA2 [64] based pipeline dadasnake, version 0.4 [65] (for settings, see Supplementary Methods S3). The consensus sequence of each SSU-ASV was aligned by BLASTn against the online database MaarjAM (accessed on 05-18-2020; [66]). SSU-ASVs with an assignment to the same virtual taxon (VT) were merged by summing up the respective read counts. All sequences of SSU-ASVs without a VT assignment were used to construct a maximum likelihood phylogenetic tree using MAFFT [67] and raxML [68]. Accordingly, SSU-ASVs in monophyletic clusters with more than 97% sequence identity were merged. The merged VTs were named in accordance with the names used in MaarjAM [66]. Clustered ASVs were named “add_cluster1+n” and sorted according to sequence abundance. In contrast, the taxonomic assignment for the ITS2 sequences was performed using the mothur implementation of the Bayesian classifier [69] and the database UNITE, version 8.2, [70] within dadasnake [65]. The tool FUNGuild, version 1.0, was used to parse fungal taxonomy and determine ecological guilds [71]. All unambiguous assignments with a confidence of “possible”, “probable”, and “highly probable” were considered. All ITS2-ASVs with an EMF-classification were considered for subsequent analyses.

### Statistical analysis

We calculated observed total AMF VT and EMF ASV richness per mixture plot by summing the unique fungal species of all tree species within a plot (same fungal species in several tree species were counted as one fungal species). We, further, calculated expected total AMF and EMF richness per mixture plot by summing up the richness of unique fungal species of the respective monocultures. Correlations between expected and observed fungal richness were tested per tree species richness level and mycorrhizal type using a linear model. The 1:1 line in the plots indicates equal expected and observed fungal richness. The observed AMF VT and EMF ASV richness, and the taxa shared between plot types were visualised using upsetR [72]. The φ (phi) specialization coefficient was calculated to determine the specialisation of each AMF VT and EMF ASV to each treatment, respectively (tree species richness x mycorrhizal type; see Supplementary Methods S4) [73]. To avoid biases due to generally rare taxa, the specialisation score of each taxon was standardised to the phi coefficients of 100 respective null models, derived by randomly swapping the mycorrhizal type of all samples. The threshold for significant specialisation was defined as 3 standard deviations from the mean of the null models.

We used linear mixed effects models to test the effects of tree species richness of the plot, tree species identity of the tree neighbour, mycorrhizal type of the target tree, and mycorrhizal type of the tree neighbour on fungal phylogenetic diversity of the target tree. We used random intercept models with plot nested in block as random factors.

To assess beta-diversity between treatments, we compared the pairwise Soerensen similarities of focal trees by extracting each pair of trees in mixed stands with its respective monoculture individual in the same block from the beta-diversity matrix (60 focal AM-tree monoculture pairs and 76 focal EM-tree monoculture pairs). In addition, all pairwise Soerensen similarities between target and neighbouring trees (35 AM-tree neighbour pairs and 39 EM-tree neighbour pairs) were extracted. The sets of similarities from each diversity level and mycorrhizal type treatment (AM-2 species, AM-4 species, EM-2 species, EM-4 species) were compared separately using Wilcoxon’s rank sum test.

## Acknowledgements

We thank the numerous helpers who supported us in the field. We thank Beatrix Schnabel, Melanie Günther, and Cynthia Albracht (UFZ) for technical assistance in sample processing and Illumina sequencing, Anja Zeuner and Romy Zeiss (iDiv) for technical assistance in determining mycorrhization rates and Kerstin Hommel (UFZ) for Sanger sequencing. The fungal composition data were computed at the High-Performance Computing (HPC) Cluster EVE, a joint effort of both the Helmholtz Centre for Environmental Research - UFZ and the German Centre for Integrative Biodiversity Research (iDiv) Halle-Jena-Leipzig, whose administrators are thanked for excellent support. This work was supported by the European Research Council (ERC) under the European Union’s Horizon 2020 research and innovation program (grant agreement no. 677232). AHB was funded by and further support came from the German Centre for Integrative Biodiversity Research (iDiv) Halle-Jena-Leipzig, funded by the German Research Foundation (FZT 118, 202548816).

