## Supplementary for "Distinct effects of host and neighbour tree identity on arbuscular and ectomycorrhizal fungi along a tree diversity gradient"

Running title: Effects of host and neighbour tree on mycorrhiza

Olga Ferlian[1,2§]*, Kezia Goldmann[3§], Nico Eisenhauer[1,2], Mika T. Tarkka[3,1], François Buscot[3,1], Anna Heintz-Buschart[1,3]

[1]German Centre for Integrative Biodiversity Research (iDiv) Halle-Jena-Leipzig, Puschstrasse 4, 04103 Leipzig, Germany

[2]Institute of Biology, Leipzig University, Puschstrasse 4, 04103 Leipzig, Germany

[3]UFZ – Helmholtz Centre for Environmental Research, Department Soil Ecology, Theodor-Lieser-Straße 4, 06120 Halle (Saale), Germany

* These authors contributed equally to this work

**Results:**

Results S1 | Sanger sequencing results for EMF

In total, the fungal associates of 109 EM root tips were successfully identified by ITS analysis, and fungi of 17 different genera were detected (Table S2). Most EM sequences belonged to Pezizales. Highest abundance (27 EM root tips) showed *Tarzetta* sequences, which were amplified from all EM trees. They mostly represented *T. catinus* and *T. cupularis* sequences. Thirteen *Peziza* sequences were obtained from *C. betulus*, *Q. petraea* and *T. platyphyllos* EM root tips, including predominantly *P. irina* and *P. ostracoderma* sequences. *Hebeloma* sequences were amplified from 14 EM root tips representing all EM trees, and they were mostly affiliated to *H. mesophaeum*. And, *Trichophaea* sequences were obtained from twelve *B. pendula*, *F. sylvaticus* and *T. platyphyllos* trees, with a predominance of *T. woolhopeia*. The fungal endophytes in the successfully amplified 34 root tips of AM trees were surprisingly diverse: they were affiliated to 22 fungal genera. Of these, five *Calyptella* sequences were amplified from *S. aucuparia* and *F. excelsior*, and four *Dactylonectria novozelandica* sequences from *P. avium, A. pseudoplatanus* and S. *aucuparia* root tips.

**Methods:**

Methods S1 | Root sampling procedure

Tree roots were followed from the tree stem to the lateral roots to ensure sampling the correct individual. Rootlets with intact fine roots were cut and stored in a plastic bag in a cooling box until further processing. To avoid contaminations between samples, tools were sterilised before taking each sample. However, twelve fine root samples could not be assigned reliably to the correct tree species and were not considered during subsequent analyses.

After sampling, the roots were immediately washed with tap water to remove attached soil, carefully swabbed, and divided into different aliquots for quantification of mycorrhizal colonisation (in 10% glycerol), individual identification of mycorrhizae (in formaldehyde), and for DNA sequencing using the Illumina MiSeq platform. Aliquots were stored at +4°C (mycorrhizal colonisation) and -20°C (individual identification and DNA sequencing), respectively, until further processing.

Methods S2 | DNA amplification

DNA concentrations were measured using a NanoDrop8000 UV-Vis spectrophotometer (Peqlab Biotechnologie GmbH, Erlangen, Germany), and all extracts were diluted to equal concentrations prior to amplification. To assess the community of AMF, we amplified the SSU following a nested PCR approach [1] using the primers GLOMERWT0-GLOMER1536 [2] and NS31-AML2 [3, 4]. To assess the community of EMF, the fungal ITS2 was amplified following the descriptions in [5] using the primers P5-5 N-ITS4 and P5-6 N-ITS4 together with P7-3 N-fITS7 and P7-4 N-fITS7 [6–8]. For both fungal targets, each sample was amplified in triplicate, accompanied by one negative control per PCR plate. The success of amplifications was checked by gel electrophoresis, triplicates were pooled together and purified with an Agencourt AMPure XP kit (Beckman Coulter, Krefeld, Germany). These cleaned products were then used as templates in a subsequent PCR, introducing the Illumina Nextera XT indices and sequencing adaptors according to the manufacturer's instructions. The amplifications had the following conditions: initial denaturation at 95°C for 3 min, eight cycles of denaturation at 98°C for 30 s, annealing at 55°C for 30 s, followed by elongation at 72°C for 30 s, and a final extension at 72°C for 5 min. Resulting PCR products were purified again with AMPure beads. Subsequently, the amplicon libraries were quantified using PicoGreen assays (Molecular Probes, Eugene, OR, USA) and pooled equimolarly. Fragment sizes and quality of DNA sequencing libraries were checked using an Agilent 2100 Bioanalyzer (Agilent Technologies, Palo Alto, CA, USA). This final pool was used for paired-end sequencing of 2×300 bp with a MiSeq Reagent kit v3 on an Illumina MiSeq platform. The sequencing was performed at the Department of Soil Ecology of the Helmholtz-Centre for Environmental Research - UFZ in Halle (Saale), Germany.

Methods S3 | Bioinformatics

Raw forward and reverse SSU (AMF) and ITS2 (EMF) reads were demultiplexed with default parameters by the Illumina reporter software v2.5.1.3 according to the index combinations, and provided as fastq files with the Illumina adaptors, indices, and sequencing primers removed. Further downstream processing was realised using the DADA2 [9] based pipeline dadasnake, version 0.4 [10]. Amplification primers were removed using cutadapt [11], allowing two mismatches for SSU reads and five mismatches for ITS reads, respectively. The quality filtering for SSU reads kept only sequences with a minimum length of 100 bp and a minimum Phred score of 20. For ITS2 reads, the minimal read length was 70 bp, also with a minimum Phred score of 20. For the merging of the sequences, a minimum overlap of 20 bp was required. After the removal of chimeric sequences, amplicon sequence variants (ASVs) were generated [12].

Methods S4 | Calculation of specialisation coefficient

The φ (phi) specialisation coefficient was calculated to determine the specialisation of each AMF VT and EMF ASV to each treatment, respectively (tree species richness x mycorrhizal type) using presence/absence data in the equation

$$\varphi = \pm \sqrt{(X^2/N)} = (a \times d - b \times c)/ \sqrt{((a + b) \times (c + d) \times (a + c) \times (b + d))},$$

with $X^2$ as the chi-square statistic for a 2 × 2 contingency table with the total number of observations N; *a* as the number of times the taxon was present in roots of trees with the respective mycorrhizal type; *b* as the number of times the taxon was present under the trees with the opposite mycorrhizal type, *c* as the number of times the taxon was absent in roots of trees with the respective mycorrhizal type; *d* as the number of times the taxon was absent under the trees with the opposite mycorrhizal type.

**Tables:**

Table S1 | Colonisation rates of arbuscular mycorrhiza and ectomycorrhiza in the ten tree species studied (intensity and frequency of mycorrhizal colonisation, respectively).

| | AM rate | EM rate |
|---|---|---|
| AM tree | | |
| *Acer pseudoplatanus* | 1.58 ± 0.84 | 0.00 ± 0.00 |
| *Aesculus hippocastanum* | 2.31 ± 1.34 | 0.00 ± 0.00 |
| *Fraxinus excelsior* | 25.81 ± 5.12 | 0.00 ± 0.00 |
| *Prunus avium* | 2.21 ± 1.30 | 0.00 ± 0.00 |
| *Sorbus aucuparia* | 1.68 ± 1.35 | 1.07 ± 1.98 |
| EM tree | | |
| *Betula pendula* | 1.52 ± 1.32 | 88.33 ± 2.04 |
| *Carpinus betulus* | 0.85 ± 0.86 | 80.49 ± 2.98 |
| *Fagus sylvatica* | 1.06 ± 0.68 | 59.49 ± 5.12 |
| *Quercus petraea* | 2.06 ± 1.59 | 74.38 ± 5.81 |
| *Tilia platyphyllos* | 1.58 ± 1.41 | 80.43 ± 3.44 |

Table S2 | Overview of mean (± SD) mycorrhizal fungal richness for all plots as well as all treatment levels.

| | AMF richness | EMF richness |
|---|---|---|
| Total | 5.63 ± 4.22 | 4.42 ± 4.20 |
| Monocultures | 4.60 ± 4.01 | 3.65 ± 3.53 |
| 2 sp. mixtures | 5.92 ± 4.57 | 4.00 ± 3.77 |
| 4 sp. mixtures | 5.65 ± 4.09 | 4.76 ± 4.49 |
| AM | 6.37 ± 5.01 | 1.36 ± 1.77 |
| AM-1 | 5.20 ± 5.14 | 0.60 ± 0.84 |
| AM-2 | 6.73 ± 5.21 | 1.47 ± 1.48 |
| AM-4 | 6.38 ± 4.95 | 1.43 ± 1.99 |
| EM | 4.88 ± 3.11 | 7.48 ± 3.65 |
| EM-1 | 4.00 ± 2.58 | 6.70 ± 2.21 |
| EM-2 | 5.10 ± 3.74 | 6.53 ± 3.66 |
| EM-4 | 4.92 ± 2.85 | 8.08 ± 3.76 |

Table S3 | Fungal taxa identified from EM root tips using Sanger sequencing. Fungal sequence distribution (represented at the genus level) is shown for the five EM tree species. Fungal taxa identified from AM root tips are not shown as we did not detect a sufficient number of sequences to be analysed.

| Host tree | Rootlets | *Cortinarius* | *Dactylonectria* | *Geopora* | *Geotrichum* | *Hebeloma* | *Helvella* | *Olpidium* | *Paxillus* | *Peziza* | *Scleroderma* | *Tarzetta* | *Trichophaea* | *Tomentella* | *Tuber* |
|---|---|---|---|---|---|---|---|---|---|---|---|---|---|---|---|
| *Betula* | 21 | 0 | 0 | 1 | 0 | 3 | 0 | 0 | 0 | 0 | 0 | 6 | 6 | 2 | 3 |
| *Carpinus* | 24 | 0 | 0 | 0 | 0 | 3 | 0 | 0 | 4 | 4 | 1 | 8 | 0 | 1 | 3 |
| *Fagus* | 21 | 0 | 0 | 0 | 0 | 4 | 0 | 0 | 0 | 1 | 3 | 6 | 3 | 0 | 4 |
| *Quercus* | 22 | 0 | 1 | 0 | 0 | 2 | 2 | 1 | 0 | 2 | 2 | 3 | 0 | 0 | 9 |
| *Tilia* | 21 | 1 | 0 | 0 | 1 | 2 | 0 | 0 | 0 | 6 | 0 | 4 | 3 | 0 | 4 |

Table S4 | Highly specialised mycorrhizal fungi with assigned fungal genus in AM and EM trees. The threshold for significant specialisation was defined as 3 standard deviations from the mean of the null models.

| AMF | Genus | EMF | Genus |
|---|---|---|---|
| **In AM trees** | | | |
| AMF01 | *Glomus* | OTU000005 | *Tarzetta* |
| AMF03 | *Glomus* | OTU000009 | *Tuber* |
| AMF04 | *Glomus* | OTU000024 | *Tuber* |
| AMF09 | *Glomus* | OTU000037 | *Hebeloma* |
| AMF11 | *Glomus* | OTU000137 | *Naucoria* |
| AMF18 | *Glomus* | | |
| AMF19 | *Paraglomus* | | |
| AMF23 | *Glomus* | | |
| AMF29 | *Glomus* | | |
| **In EM trees** | | | |
| AMF08 | *Glomus* | OTU000544 | *Pulvinula* |
| AMF12 | *Paraglomus* | | |
